# Shifts in evolutionary balance of microbial phenotypes under environmental changes

**DOI:** 10.1101/2020.03.23.003343

**Authors:** M. Kleshnina, J. C. McKerral, C. González-Tokman, J. A. Filar, J.G. Mitchell

## Abstract

Environmental conditions shape entire communities by driving microbial interactions. These interactions then find their reflection in the evolutionary outcome of microbial competition. In static, homogeneous environments a robust, or evolutionary stable, outcome in microbial communities is reachable, if it exists. However, introducing heterogeneity and time-dependence in microbial ecology leads to stochastic evolutionary outcomes determined by specific environmental changes. We utilise evolutionary game theory to provide insight into phenotypic competition in dynamic environments. We capture these effects in a perturbed evolutionary game describing microbial interactions at a phenotypic level. We show that under regular periodic environmental fluctuations a stable state that preserves dominant phenotypes is reached. However, rapid environmental shifts, especially in a cyclic interactions, can lead to critical shifts in the evolutionary balance among phenotypes. Our analysis suggests that an understanding of the robustness of the systems current state is necessary to understand when system will shift to the new equilibrium. This can be done by understanding the systems overall margin of safety, that is, what level of perturbations it can take before its equilibrium changes. In particular, the extent to which an environmental shift affects the system’s behaviour.

## Introduction

Despite the primacy of evolution in biology, there remains the critical gap in our understanding of how environment influences evolution [1]. This is especially relevant for environmental changes from the scale of small groups to global climatic events. However, the timescales for genetic adaptation are often slower than environmental changes, meaning that the important response is in the phenotype.

Bacteria are ideal model systems for studying phenotypic response due to their rapid reproduction, comparatively simple biology, and suitability for laboratory study. With most bacterial sensory systems limited to molecular uptake, it is impossible for them to anticipate environmental changes. As the most abundant life on earth, bacteria regulate the biosphere while being intimately connected to it. A single bacterial species, unlike animals, may rapidly change its role from abundant to rare, and from dominant-aggressive to rare-passive or some mix of the two depending on local habitat [2,3]. This may lead to trait variation in originally genetically identical organisms [4]. Microbial research often focuses on cooperation or competition among species, extending experimental findings to evolutionary models [5–8]. The intra-specific interactions are often overlooked. However, it is within species phenotypic variation that shapes genetic drift, which subsequently determines the genome evolution behind inter-specific interactions.

For identical genotypes in a static environments reaction to stimuli can be distinct [9]. Stochasticity in gene expression occurs in various settings and at different levels, playing a role in bacterial interactions, such as motility [10],genetic competence [11], persistence [12,13], sporulation [14], metabolism [15], and stress response [16–18]. In static environments, phenotypic robustness, rather than plasticity, is more likely to evolve, indicating that a stable phenotypic equilibrium will be reached [19]. However, under environmental change, gene expression noise may become beneficial for population fitness, indicating that stochasticity likely plays a key role in microbial communities [20]. This may extend to changing environmental conditions, where variability in fitness between the different environmental states might encourage diversification of gene expression. The lack of phenotypic robustness would then provide bacteria a better chance of managing environmental shifts, as the presence of a phenotype suited to the new conditions would confer high chances of survival. Understanding the phenotype most likely to survive is a key challenge in unravelling the complexities of the evolutionary processes within these systems.

This may extend to changing environmental conditions, where variability in fitness between the different environmental states might encourage diversification of gene expression. The lack of phenotypic robustness would then provide bacteria a better chance of managing environmental shifts, as the presence of a phenotype suited to the new conditions would confer higher chances of survival. Our definition of the lack of phenotypic robustness, or incompetency, differs from a common idea of behavioural mistakes in a way that there is no correction mechanism and that organisms will continually execute the wrong phenotype. In this context, understanding which phenotype is most likely to survive is a key challenge in unravelling the complexities of the evolutionary processes within these systems.

To investigate the properties of phenotypic interactions in evolutionary processes, we construct an evolutionary game accounting for environmental shifts. Evolutionary game theory has been widely applied to biological systems since its emergence in 1973 [21]. Analysis of microbial interactions and evolution within this setting is more recent, mostly focusing on inter-specific cooperation and competition [22–24]. However, evolutionary games are also a suitable format for examining intra-specific microbial interactions [25–29]. A significant advantage of a game theoretical approach is that it can be constructed in a cost-benefit framework, which is advantageous when studying the impacts of environmental fluctuations.

Here, we describe distinct behavioural phenotypes for bacteria. We then use an evolutionary games setting to explore how this behavioural diversity, captured as the execution of a randomised phenotype, may be beneficial (or detrimental) in different circumstances. Within this framework, we then identify which strategies are successful or unsuccessful within static or changing environments, and whether they give rise to stable or unstable systems. Finally, we provide suggestions for future theoretical and experimental work, and place the implications of our work in a broader context.

## Methods

Understanding the complex interactions within microbial communities poses a significant challenge, even before considering evolutionary processes or environmental perturbations. The ability to deconstruct interaction dynamics in this context poses computational and experimental difficulties, and possibly, impossibilities. Game theoretic approaches have the advantage of being able to elucidate the macroscale behaviour of these systems, whilst requiring few assumptions regarding the underlying drivers of emergent behaviour. In a game theoretic framework, it is possible to consider dynamic interactions between microbes where the game paradigm, reflecting environmental conditions, changes over time. Such non-steady state settings may then reveal potential shifts in the evolutionary balance within microbial communities, allowing us to suggest mechanisms for detecting which organisms survive and which become extinct.

### Modelling environmental shifts in evolution

In our framework, we consider behavioural flexibility to be the potential expression of different phenotypes when organisms are exposed to new, unfamiliar environmental conditions. As this flexibility occurs due to the imperfect realisation of the bacteria’s ‘expected’ phenotype, it is referred to as incompetence [30,31]. Incompetent bacteria may thus change their behavioural traits by switching to a different trait with some probability. The most extreme levels of incompetence are given by a limiting distribution of mistakes, captured by a stochastic matrix, *S*. Matrix *S* expresses the randomisation between different phenotypes, that is, gene stochasticity, represented here as incompetent responses. Then, the set of all probabilities that a microorganism switches its behavioural trait *i* to some other trait *j* is given by:

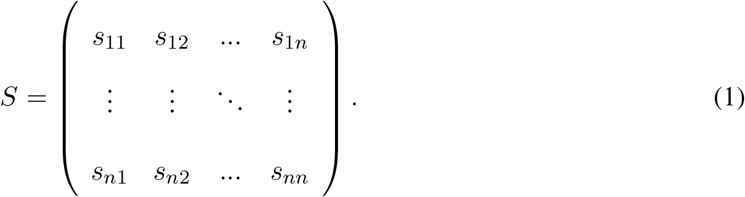

With no incompetence, matrix *S* is simply the identity matrix. Otherwise, matrix *S* reflects organisms’ responses to a new environment. If new environmental conditions are imposed on the system, matrix *I* (no mistakes, old conditions) may shift to matrix *S* (full incompetence, new conditions). We may capture these dynamics in the matrix *Q*, which describes the change in organism behaviour as they respond to environmental shifts. Matrix *Q* depends on some function λ(*τ*), reflecting the environmental dynamics, which we assume are nonlinear. Different types of changes may then be modelled as different functional forms, for example, seasonal fluctuations may be modelled using a periodic function, whereas rapid environmental shifts can be represented by an inverse sigmoid function. This implies the behavioural response to environmental change follows the dynamics:

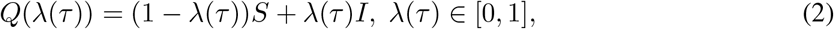

where *I* is the identity matrix and *τ* is the environmental time scale.

We assume that new environmental conditions impose new demands on organisms, which may be met by utilising an alternate phenotype. As indicated in (2), over time, there is a distribution of probabilities for the adoption of each available trait, which subsequently affects the reproduction success of different populations. This may be captured within a classic matrix-game scenario, where the fitnesses of each behavioural type are captured in the fitness matrix *R* [21]. Then, under the assumption of a changing environment, we may consider the dynamic fitness matrix as

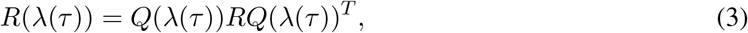

where *Q*(λ(*τ*))^*T*^ is the transpose of *Q*(λ(*τ*)). This can be interpreted in the following way. Firstly, consider pairwise interactions in a given population subjected to new environmental conditions. These organisms have a finite number, *n*, of available strategies, or phenotypes. Hence, interacting individuals compete using their chosen strategies, receiving a payoff according to the fitness matrix *R*. However, we also assume that individuals may be imperfect in their strategy execution, and may execute a different strategy according to the probabilities given in matrix *Q*(λ(*τ*)). Thus, the errors made by incompetent individuals during their interactions lead us to a perturbed payoff *r_ij_*(λ(*τ*)) from *R*(λ(*τ*)). The (*i, j*)th entry of the latter is merely the expected payoff that incorporates all possible mistakes made by the two interacting phenotypes *i* and *j*. From the evolutionary perspective, behavioural mistakes perturb population fitness over time as bacteria respond to new conditions. When the environment changes all bacteria are incompetent, except a small portion that were incompetent in the previous environment. That is, they had deficits in the old environment that are competencies in the new environment. The fitness defines reproductive success of each phenotype or behavioural strategy as

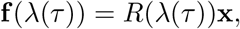

where *R*(λ(*τ*)) denotes an incompetent fitness matrix (3) with a time-varying incompetent matrix (2), and the mean fitness of the entire population is defined as follows

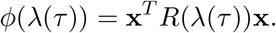

We exploit the well-studied replicator equation [32]

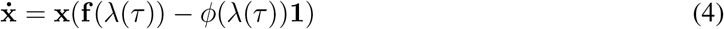

to explore the interplay between evolutionary dynamics and environmental shifts. An important feature of these dynamics is that the time scale of replicator dynamics for x(*t*) might not coincide with the time scale of environmental change for λ(*τ*). That is, environmental shifts may occur faster or slower than the interaction rates expressed by *t*. For microorganisms, however, we assume that environmental shifts happen much slower than their reproduction time scale.

### Game settings

We demonstrate the impact of environmental shifts on evolutionary balance using classic examples from evolutionary game theory: a system with an evolutionary stable state (Hawk-Dove-Retaliator game), and system with an unstable cyclic system (Rock-Paper-Scissors game). We then show how environmental conditions vary evolutionary outcomes by imposing the fluctuation dynamics λ(*τ*).

### Hawk-Dove-Retaliator

Firstly, we consider an abstract example of a stable three-dimensional system. Here, interactions between three strategies are considered: Hawk, Dove and Retaliator. This game covers a hypothetical competition between an aggressive (Hawk), passive (Dove) and mixed (Retaliator) strategies. Hawks represent an aggressive strategy, which comes with a cost of injury, whereas Doves never fight, and always share resources with a type-mates. Conversely, Retaliators will not instigate conflict but will respond to aggression with aggression, leading to conflicts arising between Hawks and Retaliators. We denote a Hawk-like strategy to characterise a hypothetical Phenotype 1, Dove strategy - Phenotype 2, and Retaliator - Phenotype 3. The fitness matrix in this case is as follows

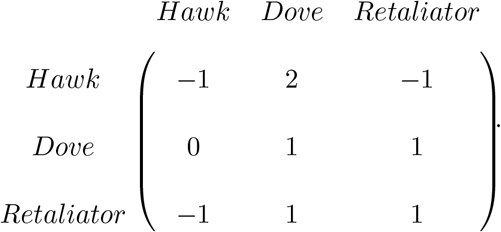

This is a widely studied game (e.g. [33]), and it has been shown that the game possesses an evolutionary stable strategy of a mixture of Hawks and Doves. However, it has also been shown that behavioural mistakes might change the evolutionary outcome [31].

### Rock-Paper-Scissors

Secondly, we examine the impacts of environmental uncertainty on a system exhibiting a cyclic relationship. The Rock-Paper-Scissors game is frequently used to describe cyclic relationships between microbial phenotypes [25–29]. It is characterised by inter-dominant strategies (Figure 3 panel a) with the following fitness matrix

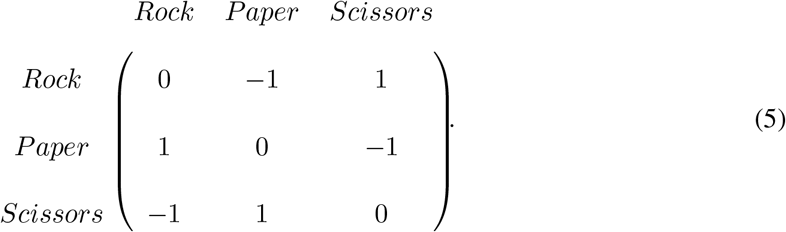

**Figure 1:**
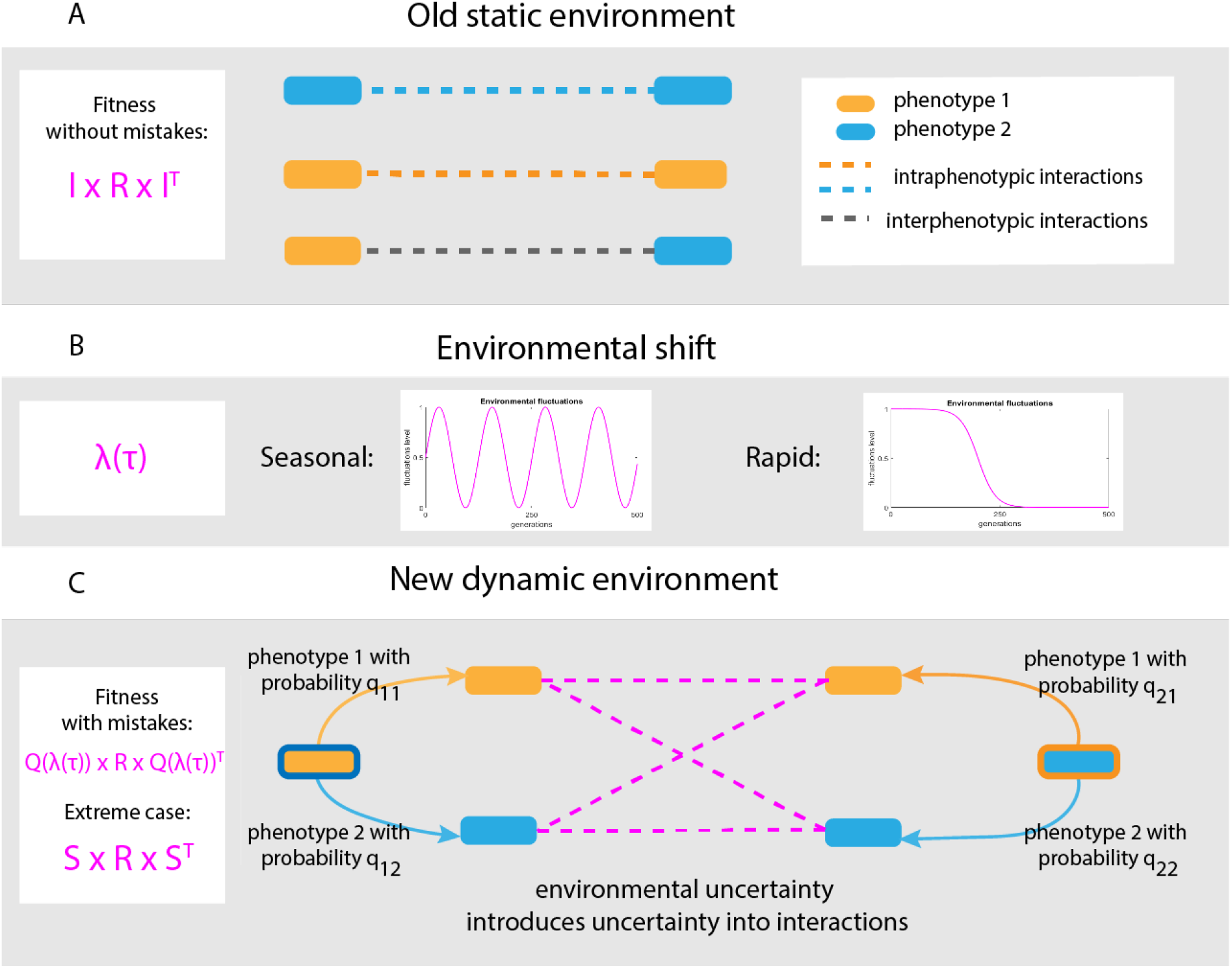
A schematic representation of the model. We assume that in static environmn-[n[['ental conditions intra- and inter-phenotypic interactions are certain and well defined. However, in new dynamic environmental conditions gene stochasticity might lead to uncertainty in interactions. In these new settings, phenotypes may switch. Thus, interactions are no longer well-defined, instead being shaped by both environmental and phenotypic expression uncertainty.

**Figure 2:**
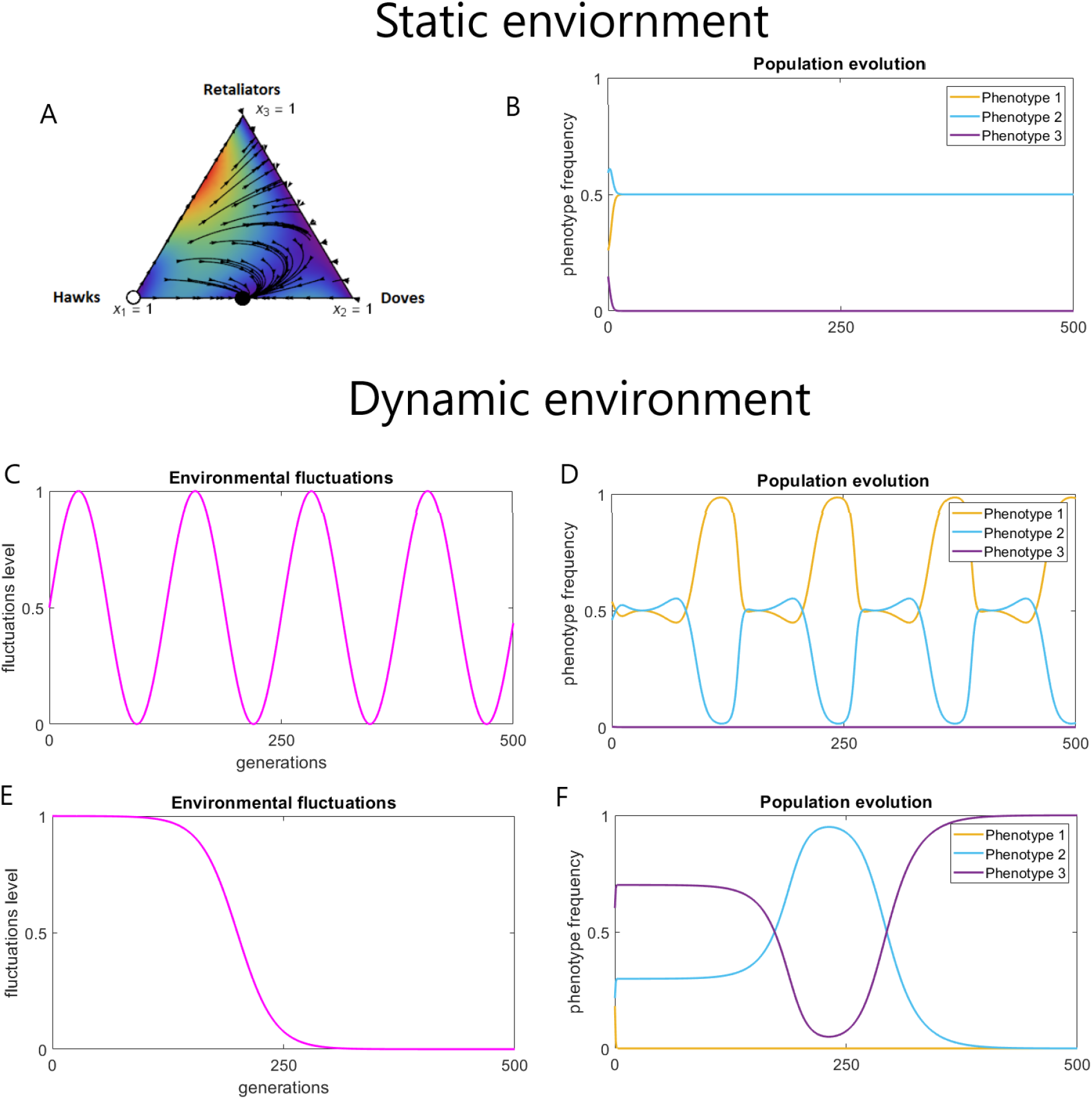
We consider an abstract example to demonstrate the interplay of phenotypic and environmental uncertainty. In static environmental conditions (panel a and b) a stable evolutionary outcome is reachable and, in this case, is a mix of Phenotypes 1 and 2. Seasonal environmental fluctuations (panels c and d) lead to a stable periodic solution, where Phenotype 1 is more beneficial. However, rapid non-periodic environmental shifts (panels e and f) can destabilise the system, leading to Phenotype 3 dominating in the new environment. This indicates that the nature of environmental changes might bring different evolutionary outcome.

**Figure 3:**
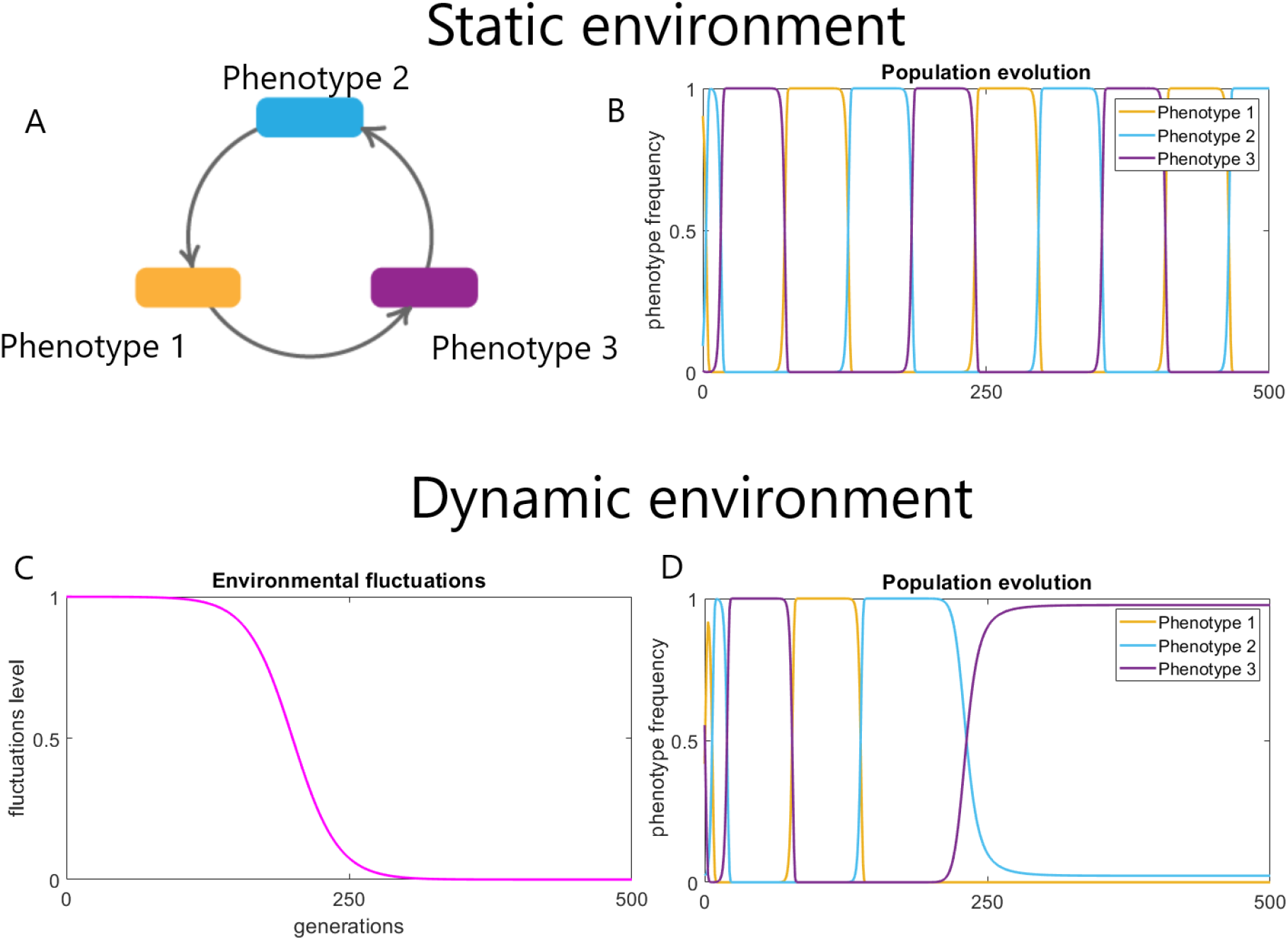
Cyclic relationships, such as Rock-Paper-Scissors, are characterised by periodic fluctuations in the phenotypic frequencies even in stable static environments (panels a and b). Under the assumptions of our model, phenotypic uncertainty together with environmental shifts destabilise the cyclic nature of phenotypic relationships, introducing a winning Phenotype (panels c and d). However, depending on the interplay between the timing of a environmental shift and the state of the system, the winning phenotype can be any one of the available phenotypes.

In this case, Phenotype 1 dominates Phenotype 3, Phenotype 3 dominates Phenotype 2 and Phenotype 2 dominates Phenotype 1. Such a relationship has been shown to exist, for example, in strains of *E.coli* [25,27]. Three different strains locally interact, resulting in an unstable cyclic system where different strains dominate at different points of time. However, this setting has only been explored under stable environmental conditions.

### Settings for modelling environmental shifts

For all simulations, from (2), we define the limiting incompetence matrix, *S*, to be

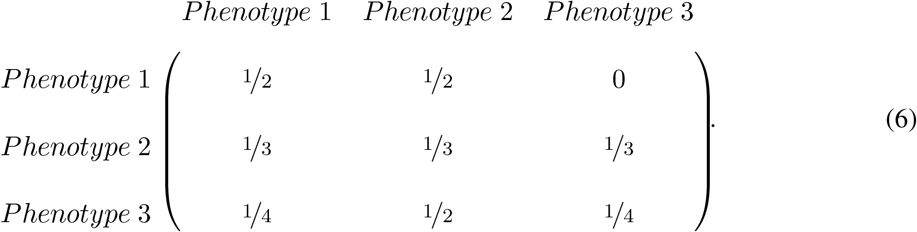

Such probability distributions provide us an opportunity to compare three abstract phenotypes which display different forms of flexibility. From (6), Phenotype 1 may exhibit Phenotype 2. Phenotype 3 is most likely to exhibit Phenotype 2 but may also express Phenotype 1, whereas Phenotype 2 is evenly random in its switching. These probabilities are taken to be high enough to reflect critical environmental shifts i.e. those which impose significant stress on the system. We chose such probabilities in order to demonstrate the effect of asymmetric probabilities distributions. In any particular case, these probabilities have to be set according to features of the population under consideration.

From (2), it is also necessary to define the environmental shift function λ(*τ*), where *τ* represents the time scale of the environmental shift. Here, we consider two functional forms. Given fast reproduction rates of microorganisms, we assume that environments change at a slower time scale than microorganisms interactions. This introduces the parameter *α*, which determines the scaling between the reproductive and environmental time scale, i.e.

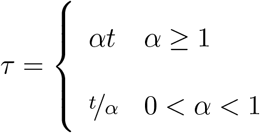

A periodic form of λ(*τ*) reflects seasonal or daily regular fluctuations, and may be defined by

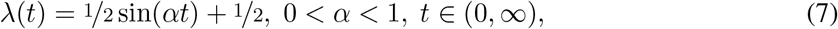

and then *α* is interpreted as a frequency of fluctuations in the environmental conditions. For instance, small *α* reflects a longer period after which the perturbation cycle begins again.

However, in the case of rapid environmental shifts, the system moves to a new, fixed state. We demonstrate this principal by extending the model to incorporate the rapid environmental shifts in the form of a reverse sigmoid function

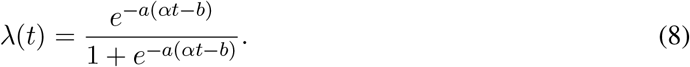

where *a* determines the steepness of the change curve and *b* is the inflection point. This function begins with a slow change rate, followed by a steep decrease before levelling of in its approach to the limiting state.

## Results and discussion

We show that such an assumption might break existing periodic relationships or lead to formation of new seasonal periodicity in the phenotypic interactions. The outcome depends on the environmental shift itself, indicating the importance of understanding how environmental changes influence affected systems.

### Periodic environmental fluctuations

The simplest setting of the Hawk-Dove-Retaliator game, with no incompetence, is characterised by existence of an evolutionary stable state [33]. This is equivalent to the outcome of interactions in a static environment. Ultimately, Hawk and Dove compete against each other and Retaliator becomes extinct (Fig. 2 panels a and b) due to its neutrality to competition.

We incorporated periodic environmental changes, described in (7), into the static model of a Hawk-Dove-Retaliator game to explore how periodicity alters evolutionary balance. Such forcing may occur through factors such as temperature changes or nutrient availability through diel or seasonal fluctuations. The adoption of alternate traits under the changed conditions may be a necessity for success or survival [34,35]. We observed that once the changes in conditions are imposed upon the system, the evolutionary outcome may change.

Periodic perturbations in the environment result in an anticorrelated relationship between populations of Phenotypes 1 and 2, following the same period of the environmental fluctuations. This indicates stability of the system is possible even in dynamic but regularly changing environments (Fig. 2 panels c and d). Indeed, we prove there exists a stable periodic solution arbitrarily close to the evolutionary stable state in a static environment (see Appendix, Periodic forcing).

Periodicity in the dominating populations can be explained from the game theoretical point of view as the organisms’ ability to react to the fluctuations through phenotypic flexibility, which becomes a key part of survival. The amplitude of the shift in populations varies depending on the rate of change in the environment and phenotype performance levels under the varying conditions. Minor shifts only bring slight change in the phenotypic frequency distributions. However, more extreme changes can destabilise the system and shift the evolutionary balance to a completely different equilibrium.

### Rapid environmental shifts

The second setting of the Hawk-Dove-Retaliator game encompasses a single, dramatic change in the environment. For example, this could be associated with an extreme weather event such as storm or flood. We apply the function (8) to the stable Hawk-Dove-Retaliator system. We shall refer to changes in dynamical behaviour caused by the changes in λ(*t*) as switching between regimes. In this case, it is not only the critical values of the parameter, but when those values are obtained that impacts the outcome of the game. Hence, we compute the critical time these switches occur. The relation between time-scales not only determines the rate of response required from organisms, but also their potential to capitalise in their interactions. Firstly, the system loses its stability at a critical time *t^c^* defined as

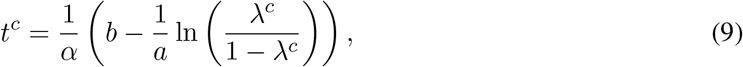

where λ^*c*^ is the bifurcation value of λ, where the dynamical system (4) changes its qualitative behaviour [31]. We show that if the environment shifts back to its original conditions, the system is also able to stabilise itself (see Appendix, Switching between regimes and critical time). Note that the most significant effect on the critical time comes from α. The importance of critical times may imply that organisms have to rapidly diversify strategies when extreme environmental conditions destabilise the system. Biologically, this would be achieved by bet-hedging, with the presence of some non-dominant phenotypes in the populations, allowing for re-emergence of those traits under some conditions [20,36–38].

For example, if *α* is small, changes may happen slowly enough for gradual adaptation. Large *α* provides a benefit to non-aggressive (low cost) phenotypes. This might happen due to an effect of cost-benefit ratio. Although high benefit often comes with higher risk, in fast changing conditions such strategies may aggressively overtake. However, in slowly changing conditions, the penalty cost becomes too expensive, meaning the non-aggressive strategies are able to succeed. In real-world settings, the laboratory can maintain strictly controlled environments, and would have a very small learning time-scaling constant *α* [39]. Conversely, marine bacteria are exposed to rapidly and dramatically changing conditions due to turbulence [40]. This means that within their bet-hedging, retaining (often) sub-optimal strategies may allow those phenotypes’ populations to explode when the high-reward strategies become favourable. Such boom-bust dynamics are frequently observed in real-world microbial systems [41].

The trajectory of environmental changes is not the only factor affecting the resulting evolutionary outcome. The construction of matrix S, or the probabilities of phenotype switching, is one of the factors determining the dominating phenotype. This may be seen in Figure 2, where the previously successful Phenotypes 1 and 2 (panel b) go extinct in a new environment (panels e and f) resulting in only Phenotype 3 surviving. Importantly, Phenotype 3 is not the most advantageous in most situations. However, in a changing environment, the phenotype may be preserved, as traits which were unfavourable before obtain a chance to persist or even overtake under new conditions. In these settings, Phenotype 3 obtains dominance due to the form of its phenotypic flexibility. This can be understood by the structure of the incompetence matrix (6), as Phenotype 1 can only randomise its phenotypic execution between Phenotypes 1 and 2, and is unable to perform Phenotype 3 at all. Secondly, Phenotype 2 randomises uniformly between all possible phenotypes, and therefore does not have a dominant executed phenotype. However, Phenotype 3 has a 50% chance of executing Phenotype 2 with other 50% being distributed equally between Phenotype 1 and Phenotype 3. This preserves all possible phenotypes in the population, but also derives an advantage under certain conditions. It may also secure its success in competition in changing environmental conditions. The capacity for phenotypic plasticity captured in this model, and our predicted scenarios under which this would be necessary, is reflected in empirical data. Static or slowly changing environmental conditions typically result in lower genomic diversity, whereas exposure to unpredictable and significant shifts promote higher diversity [27].

We shall demonstrate the effects of phenotypic flexibility further in the setting of a cyclic relationship, where the system balances out itself due to the relation between phenotypes, sometimes referred to as Rock-Paper-Scissors game.

### Shifting evolutionary balance in cyclic relationships

Next, we examine what happens under environmental shifts with a non-transitive relationship already prone to periodicity. As shown in the previous sections, the games which are inherently stable have some resilience against environmental perturbations. However, less stable systems of interactions might not survive significant environmental changes. We demonstrate this is the case utilising the example of Rock-Paper-Scissors game. This is a popular model in evolutionary game theory, whereby no strategy globally dominates or is dominated. In a stable static environment, this system exhibits cyclic fluctuations in phenotypic frequencies. In our evolutionary context, the Rock, Paper, Scissors strategies are replaced by Phenotypes 1, 2 and 3, respectively. Such periodicity has been postulated to contribute to variability in complex microbial communities [25,27]. However, this game does not posses an evolutionary stable state, meaning that any perturbation might shift the system out of its equilibrium. This is supported by the fact that non-transitive interactions are rarely observed in natural competitive communities [7].

In our model, unstable cyclic relationships must adjust to the environmental shifts induced by (8). In this case, cyclic relationships might no longer be in favour as one or another strategy may perform better under the new conditions. Naturally, one would need to determine the state of the cycle that the system was at - that is, which of the phenotypes was dominating - when the environment shifted. However, from the evolutionary perspective, this is not the only question one needs to ask. Randomised behaviour resulting in the realisation of different phenotypes can shift evolutionary balance of the system to a new equilibrium, where only one flexible phenotype will dominate (Figure 3 panels c and d). However, unlike the cases with stable systems from the previous section, in this unstable case, it is no longer possible to predict which phenotype will assume the dominating position.

As can be seen in Figure 3 panel d, the system has a limit of perturbations it can take: in the beginning of the environmental shift, the system evolves in a similar cyclic manner as in the old environment. The capacity to buffer the shift is not unlimited. Once the system hits a critical threshold, the cyclic relationship stretches its periodicity, and finally stabilises in the new state, where it will remain. In the case of such an unstable system, in order to understand which of the phenotypes will dominate, one has to examine the interplay between the critical level of environmental changes and the current state of the system. Given that the critical level of changes is hard to estimate in real microbial systems, determining the switching point becomes challenging. Nevertheless, we propose that detection of such switching points might allow prediction of how the system equilibrium will shift under certain environmental changes. For this to be done, experimental studies are required.

## Conclusions

Evolutionary and ecological theories are powerful approaches for understanding and predicting evolutionary balance of complex systems. In these complex systems stochastic gene expression is often treated as experimental noise, leading to its effects being overlooked. However, in dynamic non-stationary environments stochastic gene expression can be potentially advantageous. Here, we use two complementary models capturing periodic environmental fluctuations and rapid environmental shifts. We explored gene expression stochasticity in the framework of incompetence, where bacteria mismatch response to environmental signals. With these models we investigated the interaction of phenotypic stochasticity with environmental dynamics.

We reproduced the classical dynamics of the hawk-dove-retaliator game. In a stable environment the retaliator loses. With a single change in environment, however, the retaliator wins. Interpreting this from the microbial ecological perspective indicates that in the case of environmental uncertainty the dominant phenotype may change to one that did not initially appear evolutionarily competitive.

The inherently unstable rock-paper-scissors model is particularly appropriate for bacteria because of individuals’ metabolic flexibility and their ability to swap genes and so change their phenotype. Again, we reproduced the tripartite, alternating domination result. However, we showed that a single environmental shift can lead to one strategy or phenotype achieving continual domination.

Overall, microbes are the best theoretical and experimental models for the rock-paper-scissors game because of their inter- and intra-generational metabolic flexibility. Also, this model aptly reflects and informs microbial dynamics, where the dominant phenotype can change rapidly, leading to complex evolutionary dynamics and strategies. To understand future shifts in changing environments we need an understanding of the interaction between gene stochasticity, represented here as incompetent responses, and environmental dynamics beyond the single change response presented here.

## Appendix

The non-technical discussion presented in the main body of the text of this manuscript is based on rather detailed mathematical analyses. For the sake of completeness, in this Appendix, we outline these analyses. For more details the reader is referred to [42].

Let us first introduce some notation. Consider the replicator dynamics for the case with full competence, that is, the original evolutionary game. We shall define such a game as Γ_1_, the fully competent game^1^:

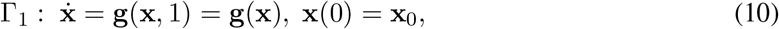

where g(x) is a replicator equation given by

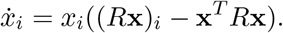

Next, we define a game with the fixed incompetence parameter as a λ-fixed incompetent game Γ_λ_:

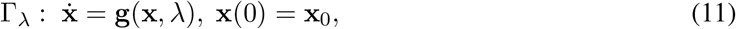

where g(x, *t*) is a replicator equation given by

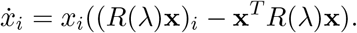

Further, we shall call a λ(*t*)-varying game Γ_λ(*t*)_ with λ being time-dependent as follows.

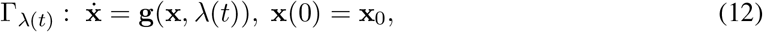

where g(x, λ(*t*)) is a replicator equation given by

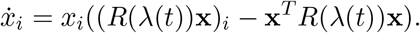

### Periodic forcing

Below, we use a result from [43] which states that a hyperbolic equilibrium solution of an autonomous differential equation persists as a periodic cycle under small periodic perturbations of the parameter. In a strict sense this statement is formulated as follows, with subscripts x and t denoting partial derivatives with respect to these arguments.

#### Theorem 1.

*[43,44] Let ẋ = G(x) be an autonomous differential equation, with* 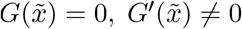, *and* 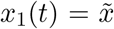 *the resulting equilibrium solution. Consider ẋ* = *G*(*x,t*) *where G*(*x,t*) *is periodic in t withperiodT. Ifforall* (*x,t*), |*G*(*x,t*) – *G*(*x*)|, |*G_x_*(*x,t*) – *G’*(*x*)|, *and* |*G_t_*(*x,t*)| *are sufficiently small, then there is a periodic solution x*_2_(*t*) *of the time-dependent equation that stays arbitrarily close to the solution of the autonomous equation*.

Now, using Theorem 1 and the fact that g(x, λ(t)) in (7) is periodic the following result can be formulated.

#### Theorem 2.

*Let* 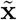 *be an ESS of* Γ_1_ *and* x_1_(*t*) *be a resulting solution for some* x_1_(0) = x_0_. *For* λ(*t*) *being a periodic function of period* 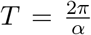 *from (7) and a sufficiently small δ, if* ||*S – I*|| < *δ, then there exists a periodic solution* x_2_(*t*) *with* x_2_(0) = x_0_ *of* Γ_λ(*t*)_ *that stays close to* x_1_(*t*).

**Proof:** In order to apply Theorem 1 we need to estimate |g(x, *t*) – g(x)|, |g_x_(x, *t*) – g’(x)|, and |g_*t*_(x, *t*)|, and show that they are sufficiently small. The first distance |g(x, *t*) – g(x)| is sufficiently small due to the Lipschitz continuity of the replicator dynamics (see p. 141 [45]), that is,

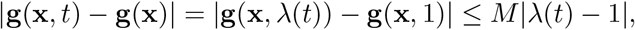

where *M* is a Lipschitz constant.

Next, we can rewrite the requirement ||*S – I*|| < *δ* as 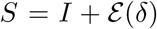, where 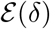 is a matrix with entries *ε_ij_*(*δ*) that are of magnitude less than *δ*. For the simplicity of notation we omit the dependence on *δ* and simply use 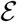. From (2) we have

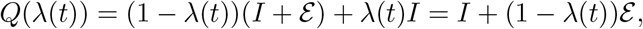

which from (3) yields

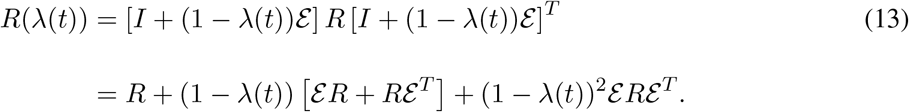

Substituting (13) into the replicator dynamics and setting *X* to be a diagonal matrix with entries of the vector x on the diagonal, we obtain

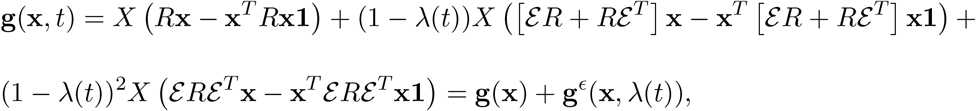

where

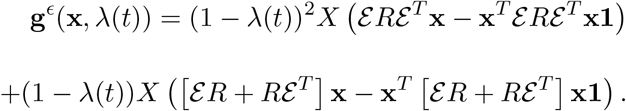

Hence,

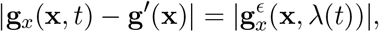

and therefore |g_x_(x, *t*) – g′(x)| → 0 as *δ* → 0.

Next, note that

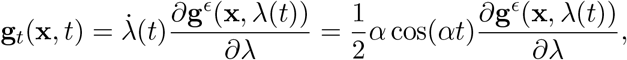

where

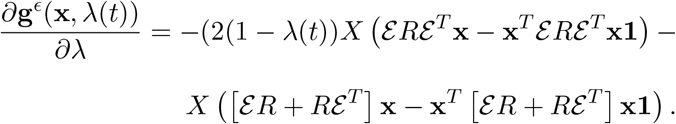

Hence, recalling once again that the entries of 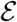 are of magnitude less than *δ*, we get that g_*t*_ (x, t) → 0 as δ → 0. The result now follows by Theorem 1.

### Switching between regimes and critical time

For this section we assume the form of the function λ(*t*) from (8). Let us recall *a maximal critical value of λ* from [31].

#### Definition 1.

*A maximal critical value of the incompetence parameter*, λ
^*u*^ ∈ [0,1], *is the maximal bifurcation parameter value*, λ^*c*^, *for the fixed point of incompetent games* Γ_λ_.

Namely, for any λ > λ^*u*^ the dynamics of Γ_λ_ remain qualitatively similar to the Γ_1_ game. Let us assume that the system Γ_λ_ has a stable equilibrium 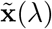 for any λ ε [λ^*u*^, 1] with a fixed point 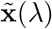 [31]. From (8) we can identify the time *t^u^* such that λ(*t^u^*) = λ^*u*^. Next, let us define the asymptotic stability of the fixed point 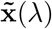.

#### Definition 2.

*Assume* λ(*t*) ∈ [λ^*u*^, 1]. *An equilibrium* 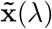 *of* Γ_λ_ *is called asymptotically stable if for any ϵ*> 0 *there exist δ*(*ϵ*) > 0, *t^ϵ^* > *t^u^ such that the solution* x(*t*) *of* Γ_λ(*t*)_ *with x*(0) = x_0_ *satisfies*

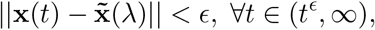

*when* 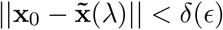.

Let us also recall Theorem 1 from [31] that the existence of an ESS in the original game implies the existence of an ESS in the incompetent game after some λ^*u*^.

#### Theorem 3.

*Let* 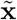 *be an ESS of the fully competent game* Γ_1_. *Let* λ^*u*^ ∈ [0,1] *be the maximal critical value of* Γ_λ_. *Suppose that* ||*Q*(λ) – 1|| ≤ *δ*(λ^*u*^), *when* λ ∈ (λ^*u*^, 1] *and δ*(λ^*u*^) *is sufficiently small, then the incompetent game* Γ_λ_ *possesses an ESS* 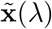 *and*

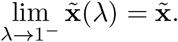

In addition, we require the next lemma based on the proof of Theorem 1 in [46].

#### Lemma 1.

*Let* 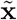 *be an ESS of the fully competent game* Γ_1_. *Then the function* 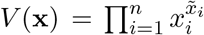 *is a strict local Lyapunov function for the replicator dynamics with the derivative along the trajectories being* 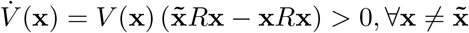, x *in a neighbourhood* 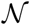 *of* 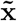.

Then the following result can now be established.

#### Theorem 4.

*Let* 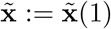 *be an ESS for* Γ_1_. Now consider the λ(*t*)-varying game Γ_λ(*t*)_, *where* λ(*t*) *is the sigmoid adaptation function (8). Let* λ^*u*^ > λ^*c*^ *for all bifurcation points* λ^*c*^ *of the λ-fixed games* Γ_λ_ *and t^u^* > 0 *be the time when* λ(*t^u^*) = λ^*u*^. *Then there exists t > t^u^ such that* 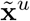 *is an ESS for* Γ_λ(*t*)_ *on the interval* (*t*, ∞) *and* 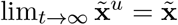.

**Proof**: An ESS is an asymptotically stable fixed point of the replicator dynamics [47]. From (2) we have

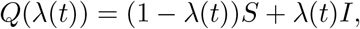

which from (3) yields

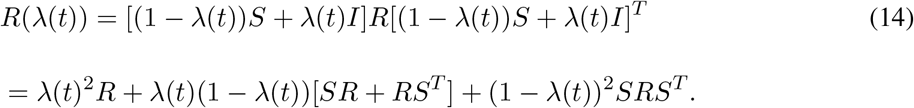

Substituting (14) into the replicator dynamics we obtain

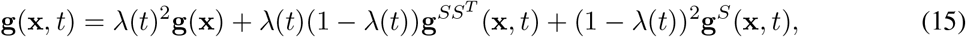

with **g**^*ssT*^ (x, *t*) = *X*([*SR* + *RS^T^*]x – x^*T*^ [*SR* + *RS^T^*]x1) and **g**^*S*^(x, *t*) = *X* (*SRS^t^*x – x^*T*^*SRS^T^*x1), where *X* is a diagonal matrix with 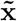 on the diagonal and 1 is a vector of ones.

Consider the function 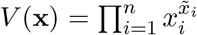 as a candidate for the strict local Lyapunov function of the equation (15). Then we have

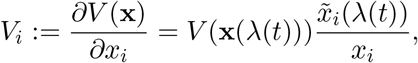

and hence from (4)

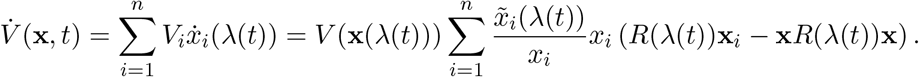

Let us denote 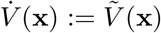. Then, substituting *R*(λ(*t*)) from (14) into 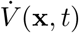 we obtain

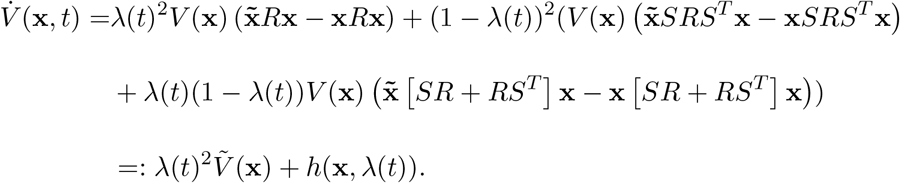

Then, as λ(*t*) → 1, *h*(x, λ(*t*)) → 0 and for every *ϵ* > 0 there exists *t^e^ > t^u^* such that ∀*t* > *t^ϵ^*

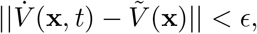

provided that x is in the attraction region 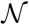 of 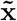 provided by Lemma 1. For every 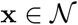 and *t* > *t^ϵ^* either 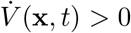 or 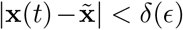 for some *δ*(*ϵ*) such that lim_*ϵ*→0_ δ(*ϵ*) = 0. As *V*(x) is a strict local Lyapunov function, then, for 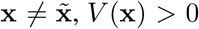. Next, let us consider the function 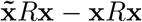, which is equal to 0 only at 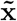 and is positive for any other x by the definition of an ESS. Then, for every *ϵ* > 0 sufficiently small there exists *δ* = *δ*(*ϵ*) such that the pre-image of x under 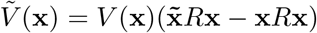 satisfies 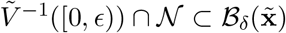 and lim_*ϵ*→0_ *δ*(*ϵ*) = 0, where 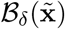 is a ball of radius *δ* centred at 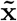. Then, 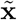 is an asymptotically stable point of g(x, *t*). By Theorem 3, as λ(*t*) → 1, the incompetent game obtains an ESS 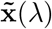 and 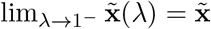.

Let us now define *the critical time* of the learning adaptation process as the first time when incompetence function, λ(*t*), attains the maximal critical value, λ^*u*^, of Theorems 3-4. Let λ^*u*^ be the maximal bifurcation value of the incompetence parameter for 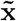, then the critical time is given by

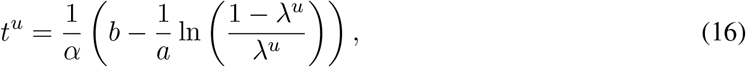

which follows immediately from the functional form of λ(*t*) given by the sigmoid function. Namely, we solve for *t^u^* the equation

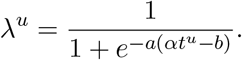

## Footnotes

1 In (10)-(12), and elsewhere, we suppress the argument, t, of differentiation in line with common practice in the field

